# Designing gene drives to limit spillover to non-target populations

**DOI:** 10.1101/680744

**Authors:** Gili Greenbaum, Marcus W. Feldman, Noah A. Rosenberg, Jaehee Kim

## Abstract

The prospect of utilizing CRISPR-based gene-drive technology for controlling populations, such as invasive and disease-vector species, has generated much excitement. However, the potential for spillovers of gene drive alleles from the target population to non-target populations — events that may be ecologically catastrophic — has raised concerns. Here, using two-population mathematical models, we investigate the possibility of limiting spillovers and impact on non-target populations by designing differential-targeting gene drives, in which the expected equilibrium gene drive allele frequencies are high in the target population but low in the non-target population. We find that achieving differential targeting is possible with certain configurations of gene drive parameters. Most of these configurations ensure differential targeting only under relatively low migration rates between target and non-target populations. Under high migration, differential targeting is possible only in a narrow region of the parameter space. When migration is increased, differential-targeting states can sharply transition to states of global fixation or global loss of the gene drive. Because fixation of the gene drive in the non-target population could severely disrupt ecosystems, we outline possible ways to avoid this outcome. Our results emphasize that, although gene drive technology is promising, understanding the potential consequences for populations other than the targets requires detailed analysis of gene-drive spillovers, and that ways to limit the unintended effects of gene drives to non-target populations should be explored prior to the application of gene drives in natural settings.

## 1 Introduction

Gene drives are genetic constructs that can bias transmission of desired alleles to progeny, allowing these alleles to rapidly increase in frequency even when they are negatively selected. Gene drives, therefore, have the potential to alter or even eradicate entire species. The population genetics of gene drives, under various genetic architectures, have been studied for several decades [1–8]. With recent innovation in technical engineering of gene drives using CRISPR/Cas9-based methods [9], gene drives have attracted considerable attention for their potential applications. In particular, engineered gene drives can conceivably alter or eradicate disease vectors, agricultural pests, or invasive species [10–14].

However, the potential of this technology also raises significant concerns due to the possibility of gene drive spillovers to non-targeted populations [10, 15–18]. The effects of such spillovers could be devastating, unintentionally driving species to extinction or permanently altering important traits, potentially leading to ecological cascades [16]. With invasive-species control, there is particular concern, because every invasive species is non-invasive in its native range. Morover, because an invasion has occurred, it is likely that invaded regions are connected to native ones through migration. As a result, to prevent gene-drive spillovers, every application of a gene drive in an invaded region must be designed to avoid them. For example, it has been suggested that CRISPR-based gene drives could be applied in New Zealand to eradicate invasive species, such as Australian possums, stoats, and rats [14, 16, 18–20]. However, such plans must account for the possibility and potential consequences of spillovers of gene drives from New Zealand to the native ranges of these species. Therefore, understanding the dynamics of gene drives with CRISPR-based constructs under migration, in the context of spillovers to non-target populations, is crucial.

Meiotic drives are systems in which Mendel’s law of equal segregation is violated, with preferential transmission of particular alleles to subsequent generations [21]. Because many general principles of population-genetic theory are violated by non-Mendelian segregation [6], models of various types of meiotic drive have been studied extensively [1–3, 5, 22–26]. Of particular interest are cases in which alleles with enhanced transmission cause reduced fitness of individuals who carry them. Analyses of meiotic drives have focused on various types of genetic architectures and evolutionary models [11], such as modifier genes [5, 25] and sex-ratio distorters [3, 26].

CRISPR-based gene drives are particular cases of preferential transmission of alleles. In CRISPR scenarios, the preferential transmission is generated at the zygote formation stage by conversion of heterozygotes carrying one copy of the gene-drive allele to homozygotes with two copies of the gene-drive allele [8, 9]. This conversion occurs when the CRISPR mechanism, which is incorporated in the gene-drive allele, edits the other chromosome to replace the wild-type allele with the gene-drive allele.

One of the features of meiotic-drive models is the existence of polymorphic equilibria, namely the states where meiotic-driver alleles persist in the population together with wild-type alleles, rather than sweeping to fixation or loss. These equilibria can be either stable or unstable, depending on the genetic architecture involved [1, 2, 4–6, 25, 27]. With CRISPR-based gene-drive technology, it has been suggested that, in a single isolated population, unstable polymorphic equilibria could be utilized for generating a biosafety measure to address the dangers of accidental releases [8]. If a gene drive is initiated at a frequency below an unstable equilibrium, it is expected to be driven to loss, whereas it is expected to be driven towards fixation if initiated at frequencies above the equilibrium [1, 8, 28–30]. Similarly, in spatially continuous populations, it has been argued that a CRISPR-based gene drive can be engineered to be driven to fixation only once it is introduced over a sufficiently large area [31].

However, few natural populations exist in isolation, and distinct populations are often connected via migration. Therefore, in order to understand potential consequences of gene-drive spillovers, explicit incorporation of migration between populations into gene-drive models is required. Some studies have examined migration models of meiotic drives with genetic architectures that have not been CRISPR-based [7, 32–37]. Among these studies, some have found that polymorphic equilibria can exist for low migration rates, but not necessarily for high migration rates.

Polymorphic equilibrium states that represent stable conditions under which gene-drive allele frequencies are high in one population but low in another might potentially be exploited to mitigate spillovers. This approach would require (1) identifying stable states in which gene-drive frequencies are high in the target population and low in the non-target population, (2) configuring the genetic architecture of a CRISPR-based gene drive to attain these states, and (3) initiating the gene drive such that it would converge to these states. We term this approach *differential targeting*. In order to consider the prospect of mitigating gene-drive spillovers through differential targeting, gene-drive models that incorporate migration and CRISPR-based genetic architectures are required.

Here, we develop and investigate such models in the context of gene-drive spillovers. We focus on identifying gene-drive designs that allow for differential targeting, and we evaluate the feasibility of the approach for mitigating spillovers.

## 2 Modeling CRISPR-based gene drives with migration

The dynamics of gene drives introduced into a wild population depend in part on features that can, at least in principle, be configured by researchers, such as the gene-drive phenotype and the conversion rate, and in part on features that cannot be controlled, such as the ecological circumstances and life-history traits of the species. In particular, the migration levels between populations are not a controlled feature of the gene drive. Here, in addition to migration, we consider several features of gene drives: (1) the selection coefficients of individuals carrying the gene-drive allele, coefficients that are related to the designed gene-drive phenotype; (2) the life-stage at which the gene-drive phenotype is expressed and subjected to natural selection, specifically whether selection acts before or after the typical migratory life stage; (3) the degree of dominance of the gene drive allele relative to the wild-type allele; (4) the efficiency of the gene-drive conversion mechanism.

We model a CRISPR-based gene drive [9], extending a previously described model [8]. We consider a population with a wild-type allele, *a*, and a gene-drive allele, *A*, which is initially absent from the population. The gene drive is characterized by (1) the conversion rate *c* of heterozygotes carrying the gene-drive allele to homozygotes of the gene-drive allele (from *Aa* to *AA*), with *c* = 1 being full conversion and *c* = 0 being regular Mendelian inheritance; (2) the degree of dominance *h* of the gene-drive allele; (3) the selection coefficient *s* of a homozygote for the gene-drive allele relative to the wild-type homozygote. In other words, the fitnesses are 1 *− s* for the *AA* genotype, 1*− hs* for the *Aa* genotype, and 1 for the *aa* genotype. For *s >* 0, the gene-drive allele is negatively selected, and we discuss only this case here, as a beneficial gene drive (*s <* 0) is expected to be driven to fixation in a deterministic model in all connected subpopulations.

### 2.1 One-deme model

We first introduce the recursion equation from [8] describing the change in gene-drive allele frequency from one generation to the next, for a single isolated population. Assuming Hardy-Weinberg equilibrium, and denoting the gene-drive allele frequency by *q*, the frequencies of the genotypes *AA*, *Aa*, and *aa* are *q*^2^, 2*q*(1 − *q*), and (1 − *q*)^2^, respectively. We denote the allele frequency in the subsequent generation by *q′*.

The *AA* homozygote fully contributes to *q′*, and is subject to a selection coefficient of 1 − *s*; in total, the relative contribution of *AA* genotypes to *q′* is *q*^2^(1 − *s*) (red arrow in Fig. 1A). Unconverted heterozygotes, *Aa*, contribute only half of the contribution of *AA*, since they pass on the *A* allele to offspring at probability 0.5. The unconverted heterozygotes represent a fraction 1 *− c* of the total heterozygotes, and they are subject to a selection coefficient of 1 *− hs*; we denote *s*_*n*_ = 0.5(1 − *c*)(1 *− hs*), and the total relative contribution of unconverted heterozygotes is 2*q*(1 − *q*)*s*_*n*_ (green arrow in Fig. 1A). Converted heterozygotes represent a proportion *c* of the heterozygotes, but because they are converted to *AA* genotypes, they fully contribute to *q′* and are subject to a selection coefficient of 1 − *s*; we denote *s*_*c*_ = *c*(1 − *s*), and the relative contribution of converted heterozygotes is 2*q*(1 − *q*)*s*_*c*_.

**Figure 1:**
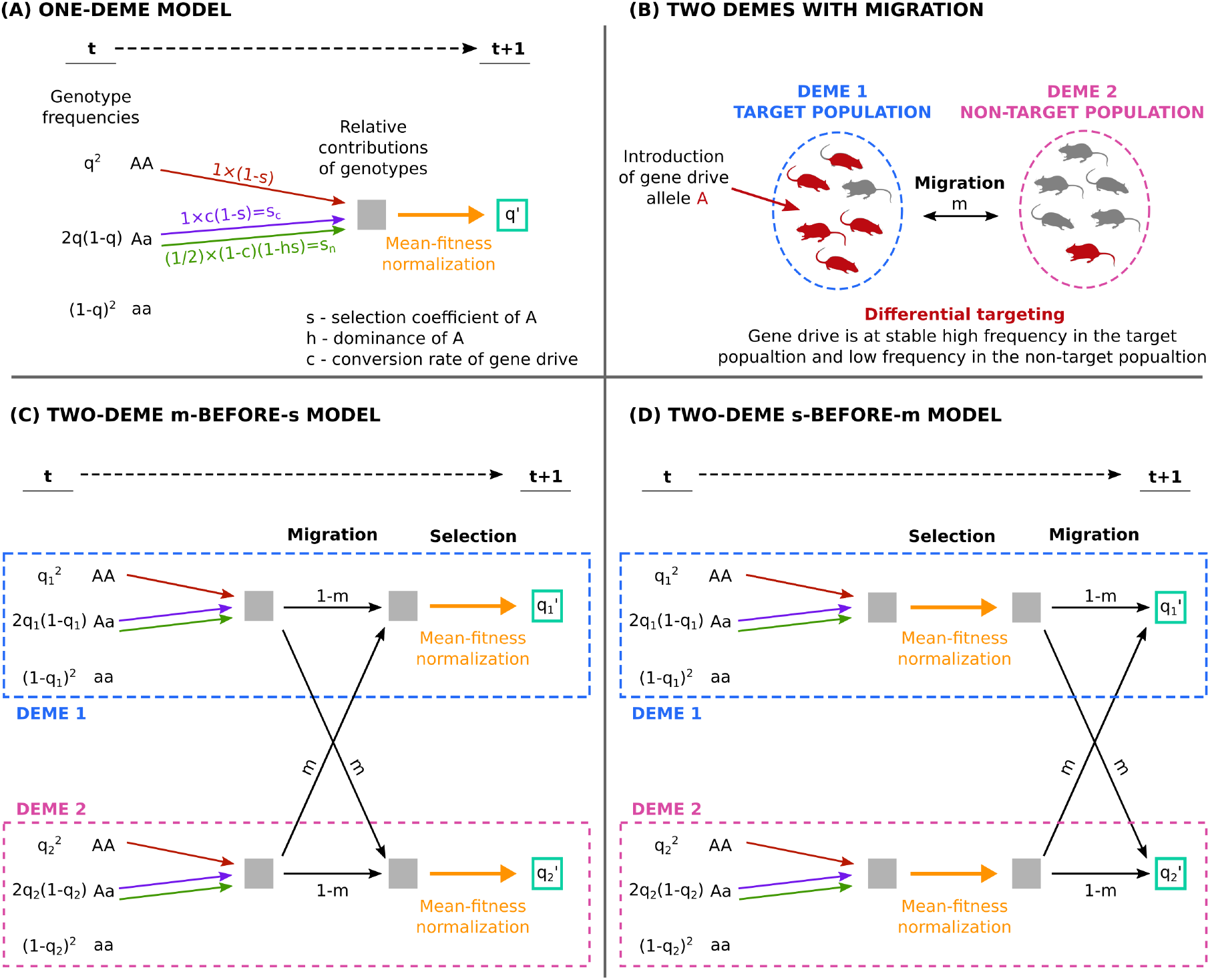
Schematic depiction of gene drive models. (A) Model of CRISPR-based gene drive in an isolated population (Eq. 1). Shown is the change of the allele frequency of the gene-drive allele A over one generation, from *t* to *t* + 1. Each genotype contributes to the frequency of A in generation *t* + 1 depending on its frequency in generation *t* and the genotype selection coefficient: the AA genotype contributes an A allele (red arrows), and the heterozygous genotype Aa contributes 1 (purple arrow) or 1/2 allelic copies (green arrow), depending on whether gene drive conversion occurs (at rate *c*). (B) A two-deme configuration with migration. The gene drive is introduced to the target population, and it can spread to the non-target population through migration. Differential targeting, when possible, would produce convergence to a stable state in which gene-drive frequencies are high in deme 1 and low in deme 2. (C) The *m*-before-*s* two-deme model (Eq. 3). Black arrows denote migration. Migration occurs before selection (Eq. 2), and 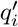 is calculated by normalizing the relative contributions to the frequency of *A* by the mean fitness of the post-migration population (orange arrow). (D) The *s*-before-*m* two-deme model (Eq. 4). Selection and mean-fitness normalization occur before migration, and 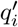 is calculated by considering the effect of migration on the post-selection allele frequencies.

Combining the three cases and normalizing by the mean fitness in the population so that frequencies in the next generation sum to 1, we obtain the following recursion relation for the gene drive allele frequency:

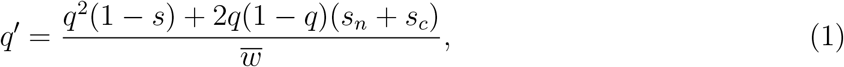

where the mean fitness is expressed as 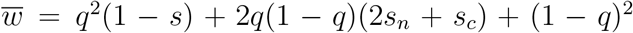. The numerator in Eq. 1 describes the genotype frequencies weighted by their relative fitnesses and their contributions to the *A* allele frequency (gray box in Fig. 1A), and the denominator normalizes this relative contribution by the mean fitness in the population (orange arrow in Fig. 1A).

### 2.2 Two-deme models

To incorporate migration between the target population and a non-target population into this modeling framework, we consider possible extensions to the one-deme model. We assume two connected demes, each large and panmictic, with symmetric migration at a rate *m* between them (Fig. 1B). As in the one-deme model, we track the dynamics of the gene drive by obtaining the change in allele frequencies of the *A* allele in the two demes, *q*_1_ and *q*_2_, respectively.

We consider two types of models that lie at two extremes for the timing of migration relative to the life stage at which the gene drive phenotype affects fitness: (1) Migration occurs before selection, meaning that individuals migrate at a relatively early life stage, and the fitness consequences of the gene-drive phenotypes are expressed in the deme to which the individuals have migrated (e.g. phenotypes expressed at late life stages, such as during reproduction). We call this the *m*-before-*s* model. (2) Selection occurs before migration, at an early life stage, and individuals experience selection in their parental deme (e.g. phenotypes expressed at the juvenile stage). We call this the *s*-before-*m* model.

#### 2.2.1 Migration before selection (*m*-before-*s*) model

In the the *m*-before-*s* model (Fig. 1C), we first consider the allele frequencies of *A* after migration but before selection, 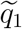 and 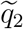 in demes 1 and 2, respectively. These frequencies are obtained by accounting for the relative contributions to the *A* allele frequencies of residents and migrants (Fig. 1C, black arrows):

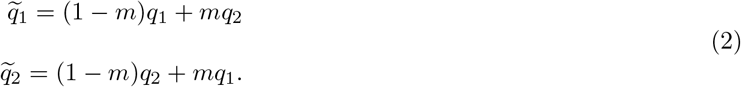

Next, we consider the frequencies of *A* after selection takes place on the post-migration gene pools (Fig. 1C, orange arrows). The changes in the allele frequencies between generations in each population are obtained as in Eq. 1, using the post-migration frequencies from Eq. 2:

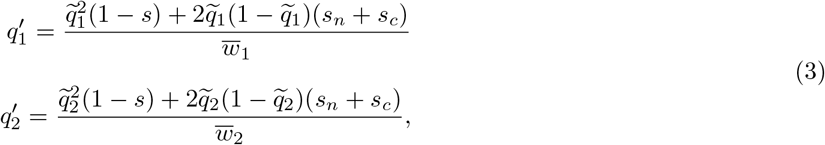

where 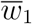 and 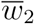 are the post-migration mean fitnesses of demes 1 and 2, respectively, expressed as 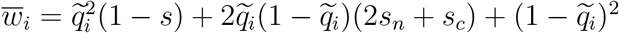.

#### 2.2.2 Selection before migration (*s*-before-*m*) model

In the *s*-before-*m* model, selection acts on the offspring in their parental deme. We determine the pre-migration frequencies of *A* by considering the relative contributions of the genotypes and normalizing by the mean fitness for each of the demes, as in Eq. 1 (Fig. 1D, orange arrows). These fitnesses, 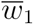 and 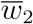, are expressed as 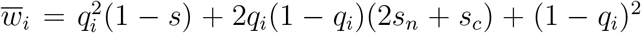. Next, we consider the impact of migration on the pre-migration gene pools of the demes, considering the relative proportion of migrants and residents, as in Eq. 2 (Fig. 1D, black arrows). Combining these two steps, we obtain:

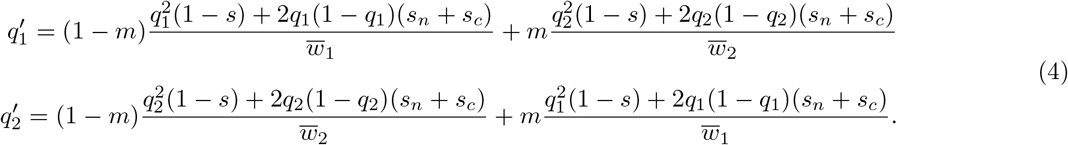

The terms on the left in the sums represent the contributions of residents to the frequencies, and the terms on the right represent the contributions of migrants.

## 3 Differential-targeting equilibria

In order to understand the evolutionary trajectories of the gene drive in the two-deme system, we study the equilibrium states of the models. This is accomplished by solving Eq. 3, for the *m*-before-*s* model, or Eq. 4, for the *s*-before-*m* model, with the equilibrium conditions 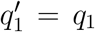 and 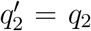 (for the equilibrium states of the one-deme model, Eq. 1, see [8]). We denote these solutions, the equilibrium points in frequency space, by 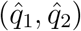, where 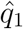 and 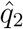 are the equilibrium frequencies in demes 1 and 2, respectively. We consider only solutions for which both 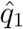 and 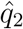 lie in the interval [0, 1], and we analyze the stability of these equilibria (see *Methods*).

A gene-drive configuration is denoted (*s, c, h*), where *s*, *c*, and *h* can have any value between 0 and 1, including 0 or 1 (i.e., the set of possible gene-drive configurations is the unit cube). Under any gene-drive configuration, there are two trivial equilibrium points, corresponding to global fixation, 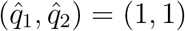, and global loss, 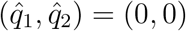, of the gene drive allele. In the one-deme model, it has been demonstrated that for some gene-drive configurations (*s, c, h*), there exists a single non-trivial (polymorphic) equilibrium point, which could be either stable or unstable [8]. For these configurations, the stabilities of the equilibria are alternating, i.e. the two trivial equilibria are stable and the non-trivial equilibrium is stable, or the two trivial equilibria are unstable and the non-trivial equilibrium is stable [8].

For the purpose of understanding how different gene-drive configurations result in different equilibrium states, we define two subsets of the set of possible configurations (*s, c, h*) (the unit cube). Both subsets have a non-trivial equilibrium in the one-deme model (Eq. 1) in addition to the two trivial ones: (1) *W*, the set of configurations for which there is an additional *non-trivial stable* equilibrium; (2) *U*, the set of configurations for which there is an additional *non-trivial unstable* equilibrium. *W* and *U* are disjoint sets, and their union is a proper subset of the unit cube because there are no non-trivial equilibrium points for some configurations (see Fig. S1). The *U* configurations are of particular interest, because they represent threshold-dependent gene drives in a one-deme system, which spread only if initiated at frequency above the unstable equilibrium [8].

We can leverage the results obtained for the one-deme model to explore equilibria in the two-deme models. For the two-deme models, the numbers of equilibrium solutions for Eqs. 3 and 4 depend on the gene drive configurations (*s, c, h*), and also on the migration rate *m* (Fig. 2). For tractability, we first consider an ecologically uninteresting yet illustrative case, where *m* = 0. In this case of no migration, the two-deme system becomes two independent one-deme systems. For *W* or *U* configurations, the number of equilibria in each one-deme system is 3, and therefore, the number of equilibria in the two disconnected demes is 3 *×* 3 = 9 for such configurations (for example, Fig. 2A). Of these equilibria, one equilibrium point, (1, 0), represents a desired outcome, in which the target deme (population 1) is affected by the gene drive, which is absent from the non-target deme (population 2). This equilibrium is stable only for *U* configurations, in each of the one-deme systems and hence in the two-deme system, and it is unstable for *W* configurations. Therefore, if a gene drive with a configuration in *U* is initiated in the basin of attraction of the stable equilibrium point (1, 0) (e.g., yellow region in Fig. 2A), then we expect that the desired outcome — the gene drive sweeping the target population and not the non-target population — will be reached.

**Figure 2:**
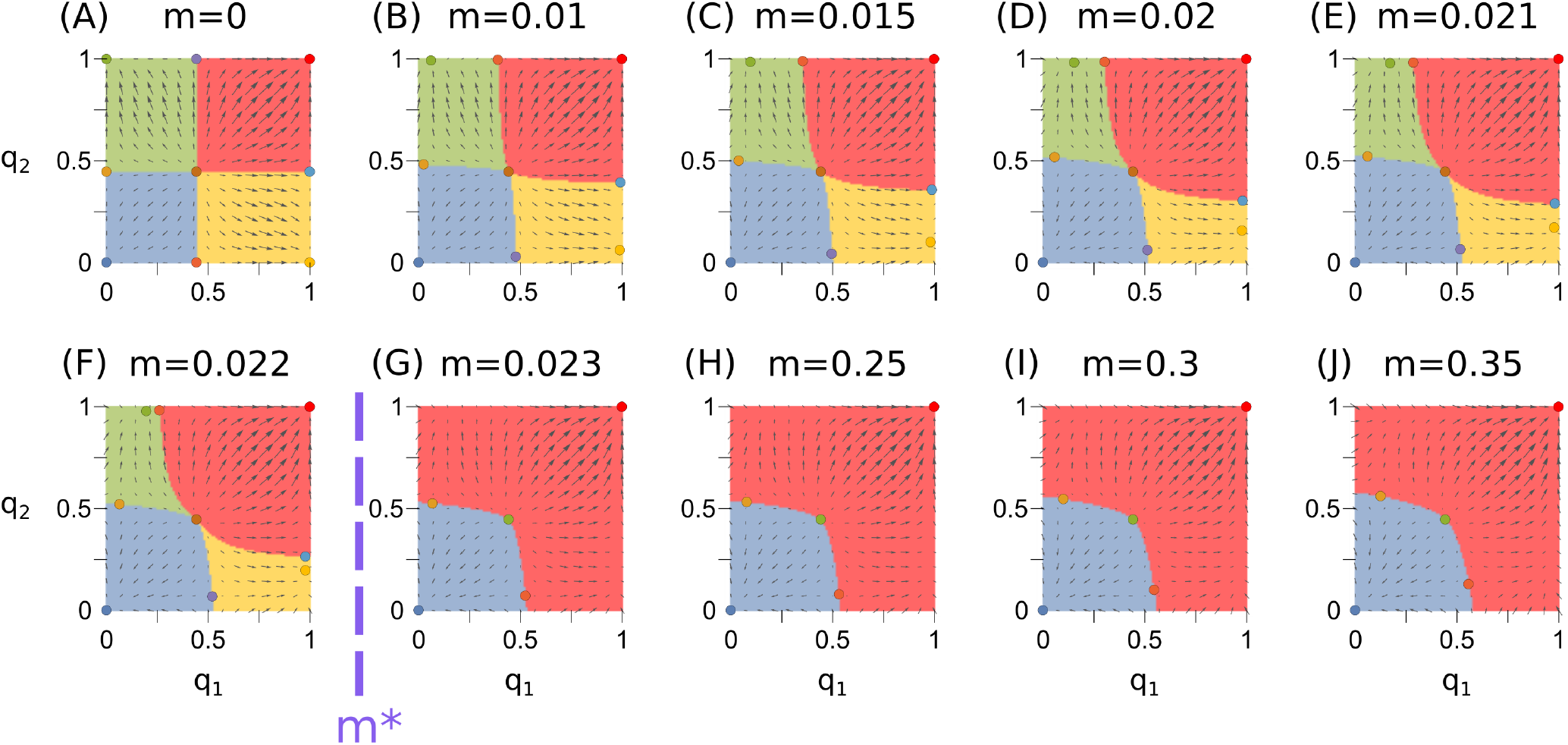
Equilibria and basins of attractions for different migration rates. Shown are results for the *m*-before-*s* model with a gene-drive configuration (*s, c, h*) of (0.6, 0.8, 0) (a configuration in *U*). The circles show the equilibria. The colored regions show the attraction basins, with the basin colors corresponding to the stable equilibria. The arrows show the vector field that describes the magnitude and direction of the change in allele frequencies at each point in frequency space. The differential-targeting equilibrium (DTE) is the stable yellow equilibrium point, which exists for migration rates lower than *m*^∗^ ≈ 0.023..

Our main interest is to understand whether similar desired outcomes, in which a stable equilibrium exists with high gene-drive frequencies in the target population and low frequencies in the non-target population, can exist for *m >* 0. We term such an equilibrium point 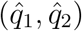, for which 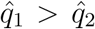, a “differential-targeting” equilibrium (DTE). In other words, we define an equilibrium point as a DTE if it is stable, and if the equilibrium gene-drive frequency in the target deme is larger than the frequency in the non-target deme. If a DTE exists, and we initiate the gene drive in its basin of attraction, then the system converges to frequencies that are higher in the target population than in the non-target population.

For *m* > 0, we explore the possible number and type of equilibria attainable in Eqs. 3 and 4 by numerically exploring the parameter space of possible gene drive configurations and migration rates. Although the gene-drive configuration sets *W* and *U* were defined for the one-deme model rather than for the two-deme models, and they are independent of *m*, they play an important role for investigating the existence of DTEs. For both the *m*-before-*s* and *s*-before-*m* models, for *U* configurations, and only for *U* configurations, we numerically find that for some *m >* 0, 9 equilibria exist. For *U* configurations, 9 equilibria exist for low *m >* 0, whereas for higher migration rates, there exist fewer equilibria (Figs. S2–S7). For example, 9 equilibria exist in Fig. 2A–F, but only 5 exist in Fig. 2G–J. When 9 equilibria exist, and only in these cases, there exists a DTE, and only one DTE. For example, in Fig. 2A–F, the yellow points are DTEs, and the yellow regions around them are the corresponding basins of attraction. At the DTE, the gene drive is not absent from the non-target population, but rather it is maintained at a frequency considerably lower than that in the target population.

Notably, for *m* > 0, we observe that DTEs exist only for low migration rates (Figs. S2–S7). For *U* configurations, when migration exceeds a certain critical threshold, we observe that there are no longer 9 equilibrium solutions to Eqs. 3 and 4, but only 5 or 3 solutions, none of which are both stable and allow differential targeting (Figs. S2–S7). We term this threshold of existence of a DTE as *m*^∗^(*s, c, h*) (Fig. 2), defined separately for Eq. 3 and Eq. 4. In other words, *m*^∗^(*s, c, h*) is defined as the supremum of the set of migration rates *m* for which there are 9 equilibria for Eq. 3 or Eq. 4 with the parameters *s*, *c*, *h*, and *m*. This set of migration rates is not empty, because for *m* = 0 there are 9 equilibria for *U* configurations, as shown above, and therefore the supremum *m*^∗^ is well-defined. For *m* > *m*^∗^, we observe that all stable equilibria are symmetric 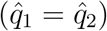. For (*s, c, h*) configurations that are not in *U*, we leave *m*^∗^(*s, c, h*) undefined, because no DTEs exist for any *m* > 0.

From the numerical solutions across the parameter space, we observe that for scenarios where *m* > *m*^∗^, only two stable equilibria exist — global fixation (1, 1) and global loss (0, 0). Hence, differential-targeting of a gene drive is not possible if *m* > *m*^∗^, and spillover to deme 2 of a gene drive that affects deme 1 is unavoidable.

## 4 Migration and differential targeting

We numerically obtained *m*^∗^ for the entire possible parameter range *U* using the numerical equilibrium solution for Eqs. 3 and 4 (Fig. 3). We find that in both models, for most of the parameter range, *m*^∗^ is low (blue regions in Fig. 3), except in a narrow curved band across the parameter space (pale yellow bands in Fig. 3). *m*^∗^ increases for higher conversion rates *c*, and it is maximal for *c* = 1 and *s* ≈ 0.7197 under both the *m*-before-*s* (*m*^∗^ ≈ 0.110) and the *s*-before-*m* (*m*^∗^ ≈ 0.109) models; we denote this maximizing selection coefficient by *s*^∗^. Note that *h* values are not relevant for a full-conversion gene drive (*c* = 1), because in such configurations, the system has no heterozygous individuals.

**Figure 3:**
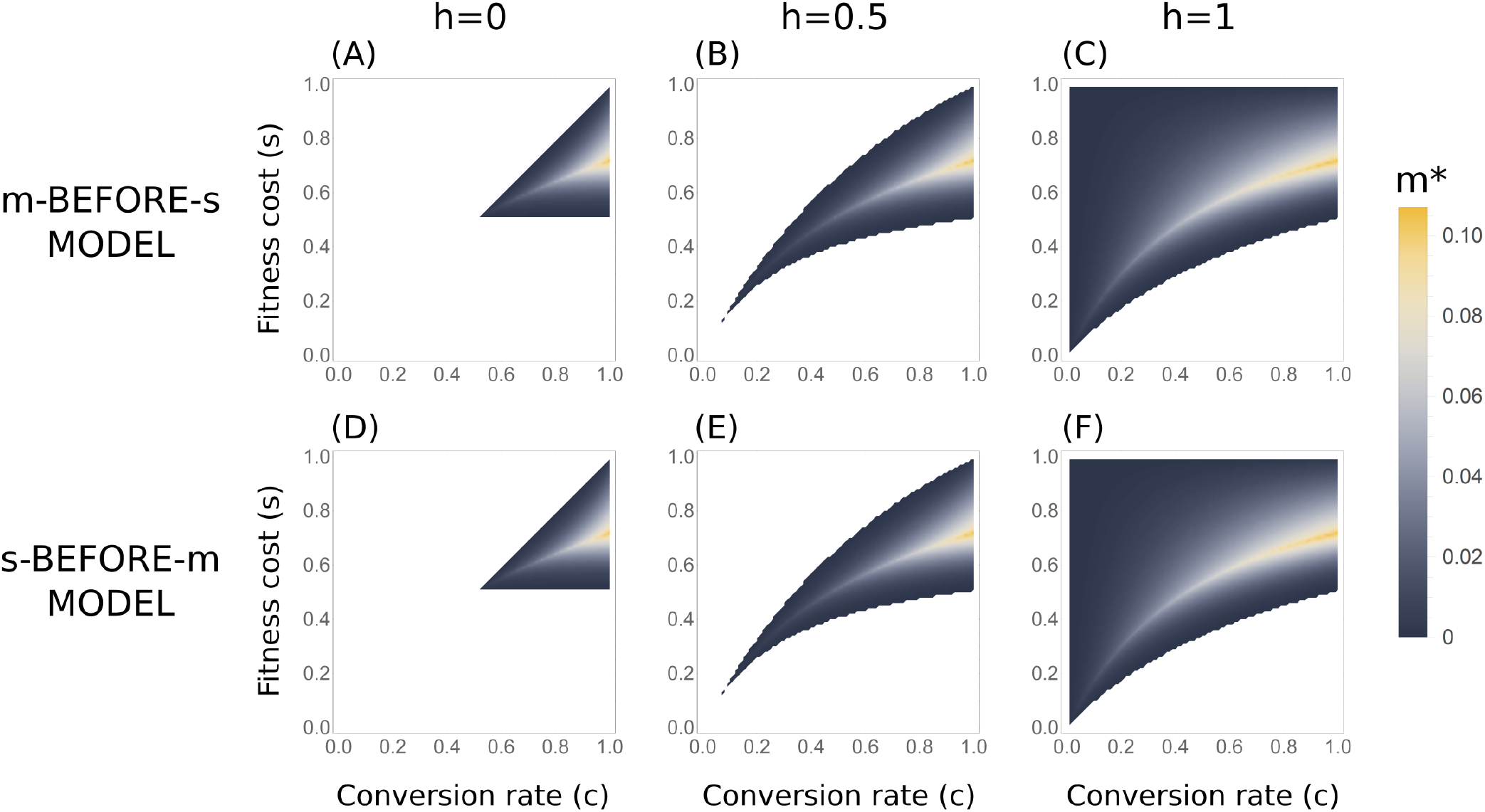
Maximal migration rates *m*^∗^ for which a differential-targeting equilibrium (DTE) exists. (A–C) *m*-before-*s* model (Eq. 3). (D–F) *s*-before-*m* model (Eq. 4). The colored regions denote the configurations for which a DTE exists (*U* configurations), and the regions in white denote configurations for which differential targeting of the demes is not possible. Over most of the parameter space, a DTE exists only for low migration rates (in blue), and only in a narrow band do DTEs exist with migration rates above *m*^∗^ = 0.05 (light yellow). In this band, *m*^∗^ values are higher for higher conversion rates, and are they are maximal, over the entire parameter space, for *c* = 1, *s* ≈ 0.7197, and any *h* value.

### 4.1 Impact of differential-targeting on the non-target population

For a gene drive with configuration (*s, c, h*) in *U* applied in a two-deme system, and assuming that, in practice, the migration rate *m* cannot be determined with certainty and is not necessarily fixed over time, it is important to understand the potential consequences for the non-target population of differential targeting, for a range of values of *m*. In principle, assuming the goal is eradication of the population in deme 1, initiation of the gene drive in the basin of attraction of the DTE leads to a high frequency of *A* in deme 1 and a low frequency in deme 2. This state is maintained until the population in deme 1 begins to collapse due to the population-level impact of the gene drive. For these generations, the population in deme 2, the non-target population, experiences the burden of the gene drive at frequency 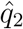 of the DTE.

For a given configuration (*s, c, h*) in *U*, we define the supremum equilibrium frequency of the DTE in the non-target population for migration rates below *m*^∗^ as 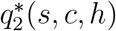. In other words, 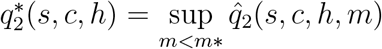, where 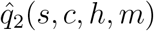 corresponds to the frequency of the DTE of a gene drive with configuration (*s, c, h*) under migration rate *m*.

We numerically computed 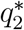 across the parameter range *U* (Fig. 4). In general, we find that 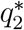 is positively correlated with *m*^∗^ (Figs. 3 and 4), meaning that gene-drive configurations that can sustain differential targeting for higher migration rates also potentially result in higher frequencies of the gene drives in the non-target population. The 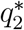 values show qualitatively similar patterns for the two models we examined, although 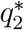 values were slightly lower in the *m*-before-*s* model than in the *s*-before-*m* model. Considering all possible gene-drive configurations in *U*, 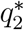 was maximal for *c* = 1 and *s* = *s*^∗^, the same configuration that maximizes *m^∗^*, in both models. Note that for a gene drive with full conversion (*c* = 1), the *h* parameter becomes irrelevant, because there are no heterozygotes. This maximal allele frequency of *A* in the non-target population at a DTE was 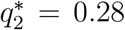 for the *m*-before-*s* model and 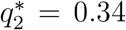 for the *s*-before-*m* model. The configurations that produce these 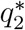 values, with *c* = 1 and *s* = *s^∗^*, delineate the conditions at which differential targeting influences the non-target population most strongly in terms of frequency of the gene-drive allele. Notably, if *c* = 1, then 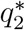 falls sharply for *s* > *s*^∗^ and less so for *s* < *s*^∗^ (Fig. 4), suggesting that deviation from *s*^∗^ is not symmetric in its impact on deme 2.

**Figure 4:**
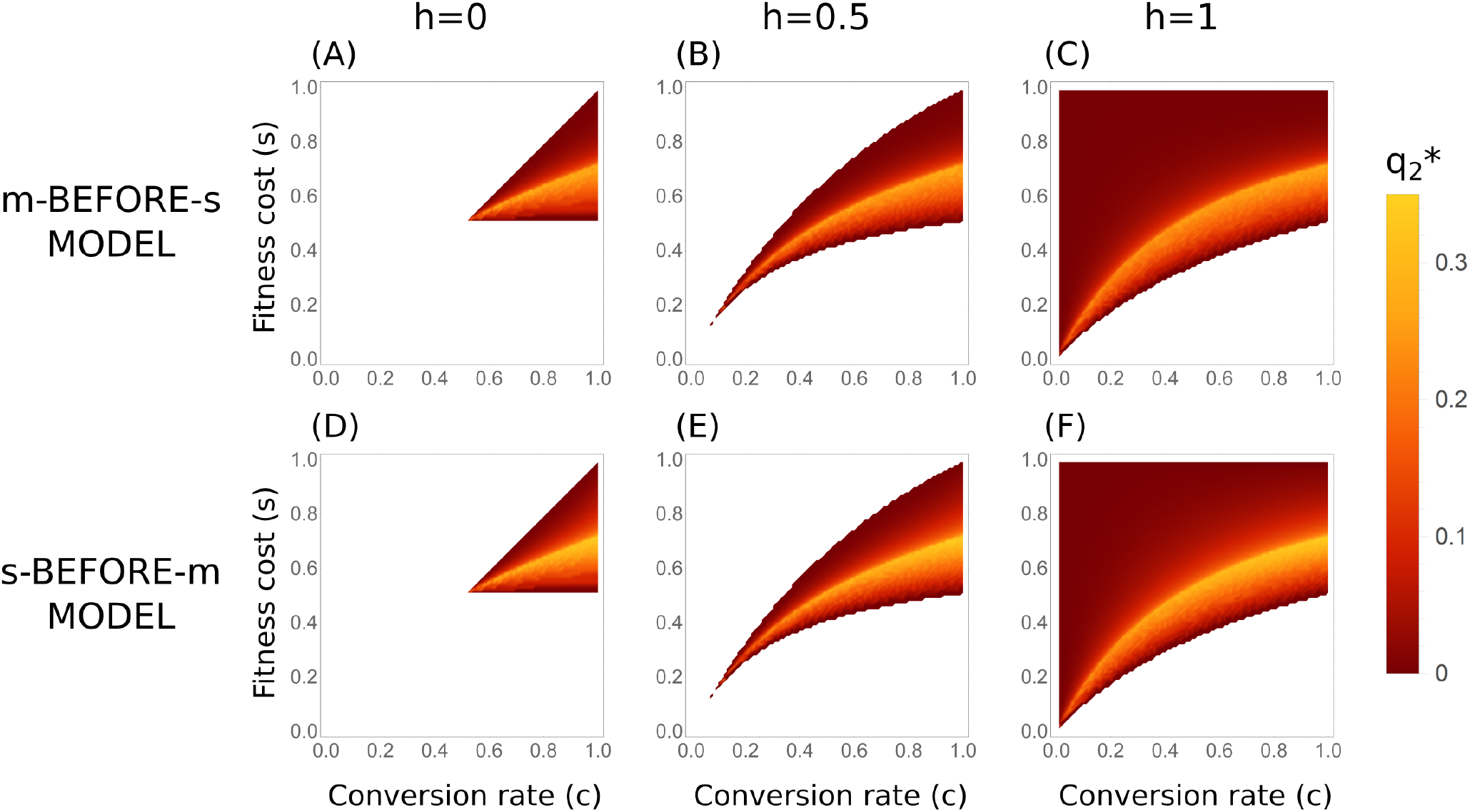
Maximal gene-drive frequencies in the non-target population, 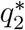, at differential-targeting equilibria (DTEs). (A–C) *m*-before-*s* model (Eq. 3). (D–F) *s*-before-*m* model (Eq. 4). Comparing to Fig. 3, 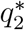 values are correlated with *m*^∗^ values.

### 4.2 Exceeding the critical migration rate *m*^∗^

For a given (*s, c, h*) configuration, the limiting migration rate at which the possibility for differential targeting can withstand migration is described by *m*^∗^. However, we must also consider the consequences of applying gene drives when migration exceeds *m*^∗^, because migration may unexpectedly change over time and may be difficult to measure. For a given configuration, if *m* > *m*^∗^, then the two-deme system is expected to rapidly converge either to global loss or to global fixation of the gene-drive allele. A gene-drive configuration that converges to global loss if *m*^∗^ is exceeded results in a failure of the gene-drive application. This case would, in general, be preferable to a gene drive that converges to global fixation when migration exceeds *m*^∗^, as the latter case would likely result in severe consequences for the non-target population.

In order to explore the outcomes for exceeding the threshold *m*^∗^, we assume that when *m* changes, the system converges rapidly to its equilibrium state, and we ignore the transient dynamics of changes between stable states. With this assumption, we can consider the continuous changes in the stable equilibrium point in frequency space due to continuous changes in the migration rate. We can track the trajectory of the DTE 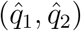 with the increase of *m* from 0 to *m*^∗^ for each gene-drive configuration (*s, c, h*) (Fig. 5). To determine the consequence of a breach of *m*^∗^, we continue this trajectory above *m*^∗^ with an instantaneous transition to either global fixation or global loss. To determine the state to which the trajectory transitions, we evaluated whether 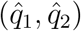, computed for the DTE with *m* = *m*^∗^ − *ϵ*, lies in the basin of attraction of global loss or of global fixation for the DTE computed for *m* = *m*^∗^ +*ϵ* (Fig. 5). For example, in Figure 2, the yellow DTE for *m* = 0.022, just below *m*^∗^, is positioned in the red global-fixation basin of attraction for the scenario *m* = 0.023, just above *m*^∗^. We therefore conclude that, for the configuration in Figure 2, the gene drive would converge to global fixation if *m*^∗^ were exceeded.

**Figure 5:**
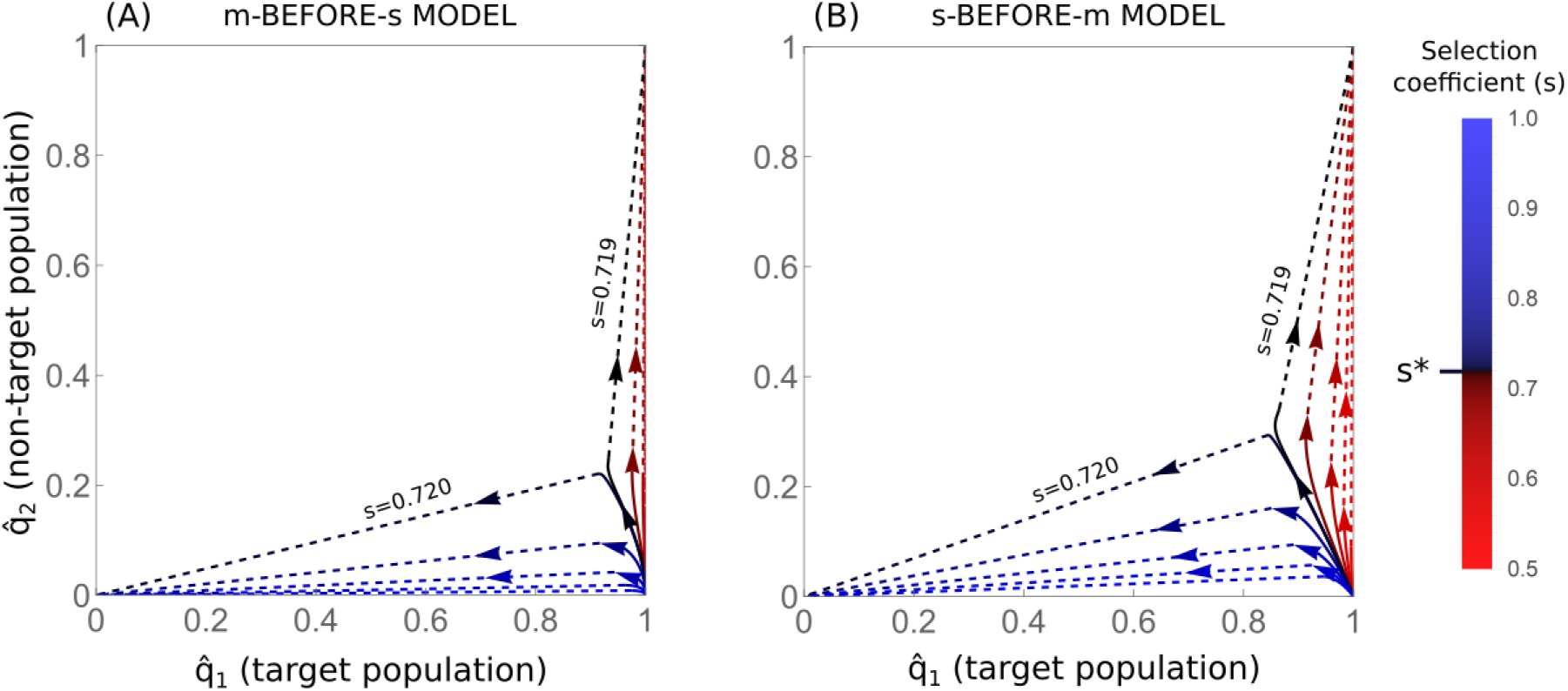
Trajectories of DTEs with increasing migration rates. Shown are trajectories for DTEs for gene drives with full conversion (*c* = 1; *h* values are not relevant in this case since there are no heterozygotes) and different selection coefficients (*s*), marked in different colors. Arrows show the direction of increased migration; solid lines show the increase of migration below *m*^∗^, and dashed lines show the abrupt transition of the equilibrium state when *m*^∗^ is exceeded, to either global fixation (1, 1) or global loss (0, 0). The color bars for *s* are centered such that the selection coefficient that maximizes *m*^∗^, *s*^∗^ ≈ 0.7197, is shown in black, selection coefficients below *s*^∗^ are in red, and those above *s*^∗^ are in blue. In both models, *s*^∗^ forms the threshold between convergence to global fixation or to global loss when the threshold *m*^∗^ is exceeded.

We computed trajectories of DTEs with increase in migration rates for *c* = 1 and different selection coefficients *s* (Fig. 5). For both models, we see that for selection coefficients lower than *s*^∗^, exceeding *m*^∗^ results in global fixation, whereas for selection coefficients higher than *s*^∗^, exceeding *m*^∗^ results in global loss (Fig. 5). In other words, for *c* = 1 and *s* < *s*^∗^, exceeding the *m*^∗^ threshold results in global fixation of the gene drive allele, whereas for *s* > *s*^∗^, a breach results in global loss of the allele. This sharp transition, and the severity of the consequences of global fixation, suggest that a safer configuration of gene drive is one with a somewhat higher selection coefficient than *s*^∗^; maintaining a safe distance from this sharp transition would be prudent, although at a cost of producing a decrease in *m*^∗^. The qualitative behavior of the system with respect to breaching *m*^∗^ is similar for *c* < 1, with a different threshold selection coefficient *s* that distinguishes global fixation from global loss.

### 4.3 Perturbations from the DTEs

So far, we have evaluated gene-drive spillovers by considering the equilibria to which the dynamics deterministically converge. However, we have not considered the possibility that stochastic events and processes might destabilize the system, allowing it to escape the DTE due to perturbations, and to reach a different stable state. In this section, we consider two types of perturbations: (1) perturbations due to genetic drift, and (2) perturbations due to an external event, such as a significant ecological disturbance, which can affect the allele frequencies of the demes.

The impact of genetic drift on the probability of escaping a DTE depends on the effective population size of each deme, *N*_*e*_. We estimated the *probability of escape* — the probability of leaving the basin of attraction of the DTE in 100 generations, starting at the DTE — by simulating the dynamics on models similar to Eqs. 3 and 4, but with allele frequencies also affected by genetic drift (see *Methods*).

To examine the impact of genetic drift, we examined the scenario of *N*_*e*_ = 100 and *c* = 1 as an example (recall that *h* has no significance for *c* = 1 because there are no heterozygotes). In general, for this scenario, the probability of escape remains below 5% until *m*^∗^ is approached (Fig. 6A–B).For larger populations, with *N*_*e*_ > 100, the probabilities of escape are expected to be lower than those in Fig. 6 due to weaker genetic drift, whereas for populations with *N*_*e*_ < 100, the probabilities of escape are expected to be higher.

**Figure 6:**
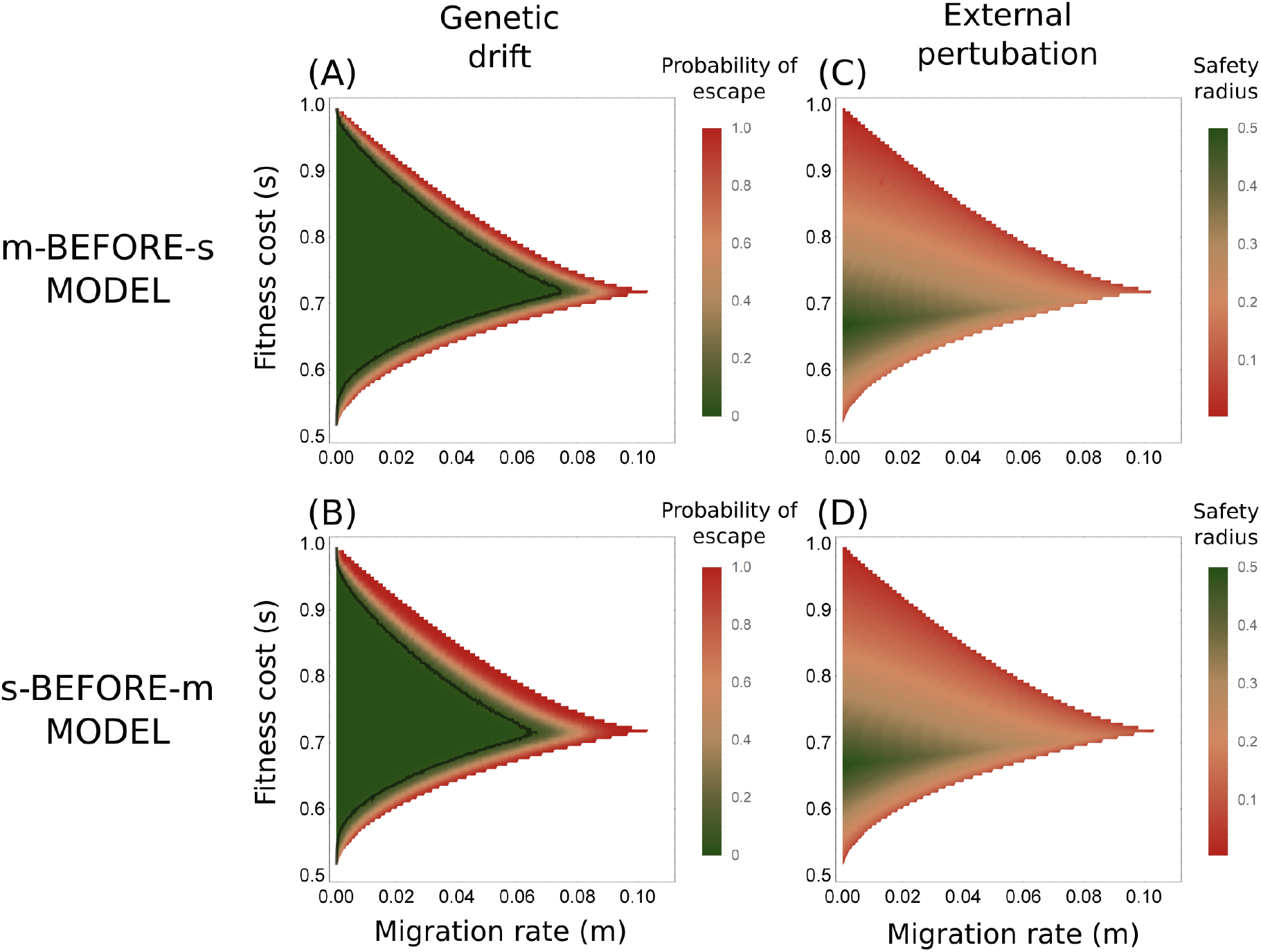
Perturbation from DTEs. Results appear for gene drives with full conversion, *c* = 1. In this case, the results are independent of *h* because there are no heterozygotes. (A) The probability of escape from the DTE due to genetic drift for different migration rates *m* and selection coefficients *s*, for the *m*-before-*s* model. The probability is defined as the probability of departing from the attraction basin of the DTE over 100 generations with genetic drift, in a Wright-Fisher population with *N*_*e*_ = 100. The black line denotes 5% probability of escape. (B) The probability of escape for the *s*-before-*m* model. (C) The safety radius of the DTE for the *m*-before-*s* model. (D) The safety radius of the DTE for the *s*-before-*m* model. In (A) and (B), probabilities were estimated with 1000 simulated replicates. In all panels, white regions denote scenarios for which *m* > *m*^∗^.

Genetic drift is a source of perturbation that is internal to the system, but we also consider external perturbations that alter allele frequencies in the demes. In this case, we evaluate the resilience to perturbations by examining the consequence of a single perturbation to the eventual DTE of the system. We define the safety radius of a gene-drive configuration under a given migration rate to be the maximal magnitude of a perturbation from a DTE, in the allele frequency space, above which the system will not converge back to the DTE. In other words, the safety radius is defined as the shortest Euclidean distance from the DTE to the boundary of its basin of attraction. For perturbations from the DTE larger than the safety radius, the system is expected to converge to an equilibrium that is not a DTE. Similar to the probability of escape from the DTE by genetic drift, the safety radii are particularly small when migration rates approach *m*^∗^ (Fig. 6C–D).

## 5 Discussion

We have explored the possibility of effectively applying a gene drive in a target population while limiting the exposure of a non-target population to spillovers. By investigating equilibria of the expected evolutionary dynamics, we have shown that for some gene-drive configurations (*s, c, h*), those for which a polymorphic unstable equilibrium exists in the one-deme model (U configurations), it is possible to initiate the gene drive such that differential targeting — higher equilibrium gene-drive allele frequencies in the target population than in the non-target population — is possible. However, we have also shown that for these configurations, upon increasing the migration rate *m*, a sharp transition occurs to a state in which the differential-targeting equilibrium no longer exists, and the two populations face similar fates, either global loss or global fixation of the gene-drive allele. We numerically identified an “optimal” gene-drive configuration, with *c* = 1 and *s*^∗^ ≈ 0.7197, as the configuration that allows for differential targeting under the highest migration rate, *m*^∗^ ≈ 0.110.

For most *U* configurations, other than in a narrow region of the parameter space for (*s, c, h*), differential targeting can be achieved only for low migration rates (Fig. 3). Consequently, for relatively high migration rates, it is unlikely that gene drives could be configured with enough accuracy to provide sufficient confidence that differential targeting could be achieved. In general, under the models, gene drive spillovers to the non-target population should therefore be considered likely, and prevention of spillovers should in most cases not rely on differential targeting. Only when the gene-drive configuration (*s, c, h*) has been validated with high accuracy, or when low migration rates between target and non-target populations can be assured, will it be possible to view differential targeting as a practical measure for limiting spillovers.

We found that the configuration that maximizes the critical migration threshold *m*^∗^ (*s*^∗^ ≈ 0.7197, *c* = 1, and any *h* value) is also the one that maximizes the impact on the non-target population in the sense of producing the largest *q*_2_^*∗*^ (Fig. 4). Allowing the selection coefficient to vary, this same configuration also differentiates between scenarios for which increasing migration above *m*^∗^ results in global loss or global fixation of the gene drive allele (Fig. 5). In practical settings, such breaches may occur as a result of unexpected increases in migration, or from errors in approximation of actual migration rates. As a precaution, it would be wise to design gene drives (*s, c, h*) for which exceeding *m*^∗^ results in global loss rather than the alternative, in order to avoid full exposure of the non-target population to the gene drive.

In this regard, considering practical difficulties of accurately specifying gene-drive parameters such as *s*, *c*, and *h*, it is advisable to avoid gene-drive configurations that are too close to this sharp transition of *s*^∗^, and to design *s* values that are somewhat higher than *s*^∗^. This strategy would also result in lower impact on the non-target population in terms of gene-drive allele frequencies 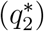, compared to values that are slightly lower than *s^∗^* (Fig. 4), albeit at the cost of decreasing *m*^∗^ (Fig. 3).

Although some minor differences exist between the outcomes for the two models we examined, in general, the order of migration and selection has little effect on the properties studied here. The migration thresholds were almost identical in the two models, and the maximal migration threshold was negligibly higher in the *m*-before-*s* model (0.110 vs. 0.109; Fig. 3), and was attained for the same gene drive configuration in both models (*c* = 1 and *s* = *s*^∗^). The main difference between the models was in the maximal DTE allele frequency of the gene drive in the non-target population, 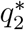, which was slightly higher in the *s*-before-*m* model (Fig. 4).

For organisms inhabiting a large geographic range, it is important to identify the relevant scale to which the migration rates *m*, which are defined as migration probabilities of individuals, apply. These rates refer to populations in the region from which migration occurs. For example, considering a target island with an invasive rodent species and a non-target island with native rodents, and assuming that the rodents migrate mostly via major ports on these islands, the subpopulation of a port and its immediate surroundings, and not the population of the entire island, would likely be the appropriate unit for estimating *m* and *N*_*e*_. Migration rates among subpopulations on an island are typically much higher than migration rates between populations on different islands. Therefore, for migration between two highly-populated islands, for example, if we measure *m* using the entire island population, and not among appropriate subpopulations, then we might significantly underestimate the appropriate *m* for the purpose of our models.

Before CRISPR-based gene drives can be efficiently and safely applied, many challenges, such as the potential evolution of resistance to the gene drive modification [38–44], must be overcome. For the type of differential targeting described here, an accurate configuration of gene drive parameters, *s*, *c*, and *h*, is required. However, it is unclear whether such accuracy can be feasibly obtained. For example, to date, conversion rates *c >* 90% have been demonstrated only in some invertebrates [9, 45–48] and yeast [49], with much lower conversion rates occurring in mammals [50]. Another difficulty for differential targeting is that implementation might require a large concentrated effort to initiate the gene drive in the basin of attraction of the desired equilibrium (the DTE). This effort could require high gene-drive allele frequencies, and hence it might involve engineering and releasing a very large number of engineered individuals. A third problem is that we do not yet understand the consequences of more complex population structure, involving more than two populations or populations that are internally structured. Given these difficulties, to understand and mitigate the dangers of spillovers, it will be important to study more elaborate population structures in models of gene drives, as well as to explore additional biosafety mechanisms and molecular safe-guarding techniques [37, 51–56].

CRISPR-based gene-drive technology is potentially a powerful tool for species control, but it also has many associated risks. That scenarios exist that permit differential targeting should not be read as a call for applications of gene drives in wild settings, or as a blueprint for CRISPR-based gene-drive design. Instead, we believe the consequences of gene-drive application and the potential for spillovers should be further discussed in the scientific community and in society at large before actual gene-drive applications are realized, and a goal of this paper is to facilitate and inform these discussions.

## 6 Methods

### 6.1 Equilibria

For a given model (*m*-before-*s* or *s*-before-*m*), gene-drive configuration (*s, c, h*), and migration rate *m*, the equilibria were computed by numerically solving the systems of equations in Eq. 3 or 4, with the equilibrium conditions 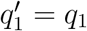 and 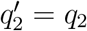, using Mathematica software [57].

### 6.2 Stability of equilibria

For each equilibrium 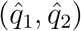, its stability was determined by examining how the trajectories of the allele frequencies in the two populations behave in its vicinity in allele frequency space.

The system of discrete-time recursion equations that describe the change of allele frequencies over a single generation are:

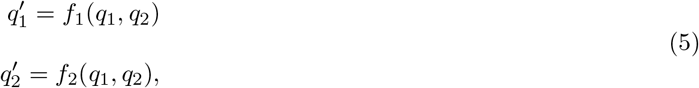

where *f*_*i*_ (for *i* = 1, 2) is defined in Eq. 3 or 4, depending on the model. To analyze the stability of the system in the vicinity of an equilibrium 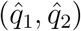, we linearized the nonlinear functions *f*_1_ and *f*_2_ by taking the first-order approximation of the Taylor expansion about the equilibrium. From this linearization, in order to understand the consequences of small perturbations from the equilibrium, we examined the matrix of the partial derivatives of *f*_1_ and *f*_2_, evaluated at the equilibrium (i.e., the Jacobian matrix of Eq. 5):

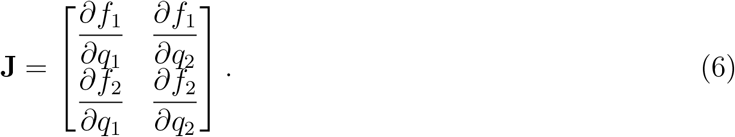

The stability type of the equilibrium point 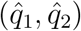 was determined by examining the two eigenvalues *λ*_1_ and *λ*_2_ of **J** evaluated at 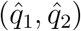. If both eigenvalues have modulus less than 1 (*|λ*_1_*| <* 1 and *|λ*_2_*| <* 1), then the equilibrium point 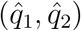 is stable. If at least one eigenvalue has modulus greater than 1, then the equilibrium point is unstable [58]. For the special case with *|λ*_1_*|* = 1 or *|λ*_2_*|* = 1, see [58].

To generate Fig. 2 and Figs. S2–S7, we computed the equilibria and determined their stability for all (*s, c, h*) combinations, with *s* ranging from 0 to 1 in increments of 0.01, *c* ranging from 0 to 1 in increments of 0.01, and *h* = 0, 0.5 or 1. To generate Figs. 3 and 4, these configurations were examined for each migration rate *m* between 0 and 0.15 at increments of 0.0001.

For a stable equilibrium, there exists a basin of attraction in allele frequency space around the equilibrium [59], such that any trajectory starting within the basin eventually converges to the stable equilibrium point. To determine basins of attraction for a configuration (*s, c, h*) and migration rate *m*, we start from each point (*q*_1_*, q*_2_) in the frequency space, and track its trajectory for 1000 generations by iteratively using Eq. 3 or 4. We denote the endpoint of the trajectory with initial condition (*q*_1_*, q*_2_) by (*q*_1_*, q*_2_)_1000_. The attractor associated with (*q*_1_*, q*_2_) was defined as the equilibrium 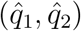 that was at a Euclidean distance less than 0.001 in frequency space from (*q*_1_*, q*_2_)_1000_. If there were no such equilibria or more than one such equilibrium, then the attractor of (*q*_1_*, q*_2_) was left undefined. The basins of attraction were computed for (*q*_1_*, q*_2_) points at a resolution of 0.01 *×* 0.01 in frequency space. For all computations used to generate Figs. 2, 5, and 6, attractors were defined for all points computed.

### 6.3 Genetic drift

To determine the probability of escape from DTEs, we formulated the models as a stochastic process. We modified the deterministic equations in Eqs. 3 and 4 such that the frequency in each generation is binomially distributed with 2*N*_*e*_ trials, following the Wright-Fisher model for diploid populations, for each population, with the same means as for Eqs. 3 and 4. Genetic drift was implemented after both selection and migration, in both models.

The distribution of the gene-drive allele frequency following one generation in the *m*-before-*s* model is given by:

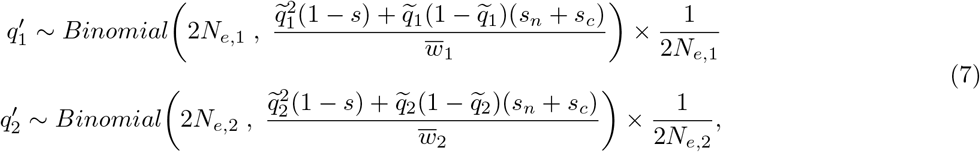

where *N*_*e*,1_ and *N*_*e*,2_ are the effective population sizes in demes 1 and 2, respectively.

Similarly, for the *s*-before-*m* model, the equations describing the change in frequency of *A* over one generation are:

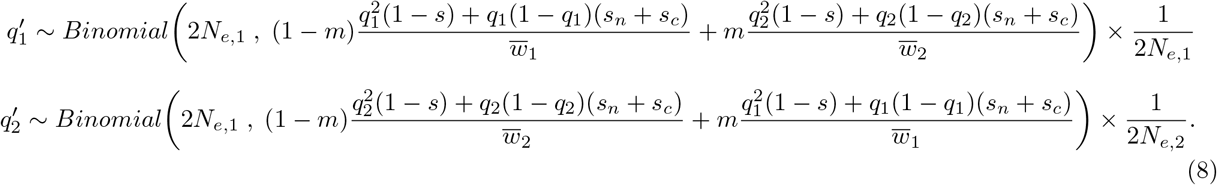

With these models, we can simulate the allele frequency trajectories in the two demes, accounting for the effective population sizes. For Fig. 6A and 6B, we generated trajectories for *N*_*e*,1_ = *N*_*e*,2_ = 100 starting at the DTE allele frequencies. We therefore consider the probability of escape once the DTE is reached, and not the probability of escape during the transient convergence to the stable state when the gene drive is initiated at allele frequencies other than the DTE frequencies.

For each gene-drive configuration (*s, c, h*) and migration rate *m*, the equilibria and basins of attraction were determined. Next, for each (*s, c, h, m*), 1000 simulations of the stochastic process were initiated, starting at the DTE 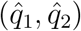, and they were iterated for 100 generations by repeatedly applying Eq. S1 or Eq. S2. In each simulation, it was determined whether or not the allele frequencies remained in the basin of attraction of the DTE during all 100 generations. The probability of escape was computed as the fraction of the simulations for which the allele frequencies did not remain in the attraction basin of the DTE. Trajectories that escaped the basin of attraction at any time were considered as escaped trajectories, irrespective of whether or not they returned to the basin of attraction, so as to provide a conservative estimate of probability of escape.

### 6.4 Safety radius

In order to compute the safety radius for a given model, gene-drive configuration, and migration rate, the equilibria and basins of attraction were first computed. The Euclidean distances, in frequency space, between the DTE and all points in its basin of attraction were then computed. The safety radius was defined as the infimum over distances, in frequency space, between the DTE and points outside its basin of attraction. In other words, for a DTE 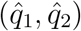 with basin of attraction 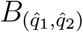, the safety radius was defined as 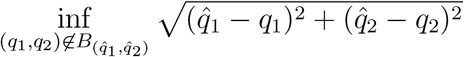. In practice, the safety radius was determined by computing the minimal distance between the DTE and points outside the basin of attraction, for points at 0.01 *×* 0.01 resolution in frequency space.

## Appendix A Characterization of equilibria

**Figure S1:**
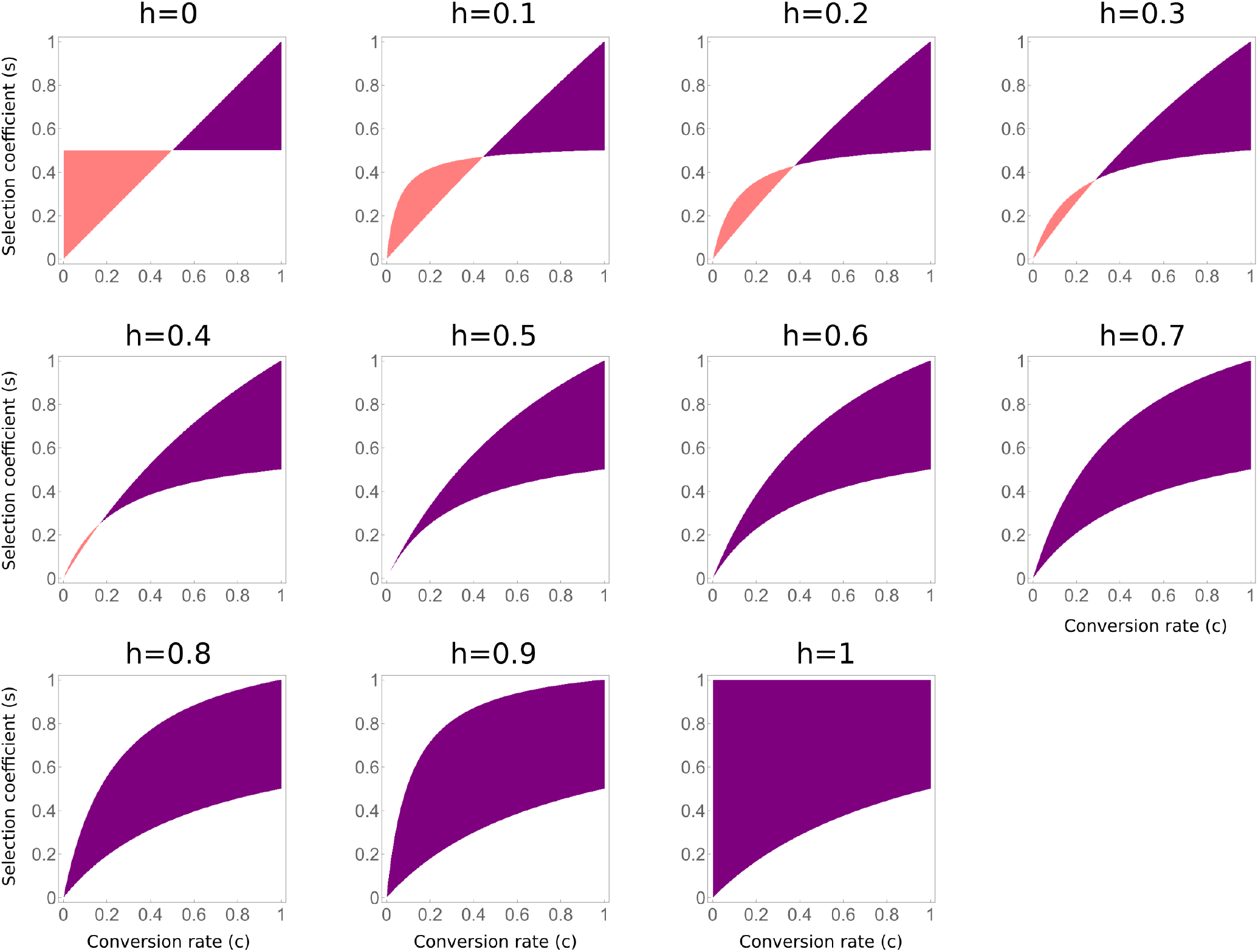
Gene drive configurations with non-trivial equilibria in the one-deme model (Eq. 1). Purple — *U* configurations, with 1 unstable non-trivial equilibrium, and 2 stable trivial equilibria (fixation 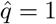, and loss 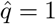); pink — *W* configurations, with 1 stable non-trivial equilibrium, and 2 unstable trivial equilibria (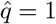 and 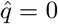). In white are configurations for which there are only 2 trivial equilibria, and no non-trivial equilibria. The panels show results for 11 values of *h*, and all possible values of *s* and *c*. See [8] for further details.

**Figure S2:**
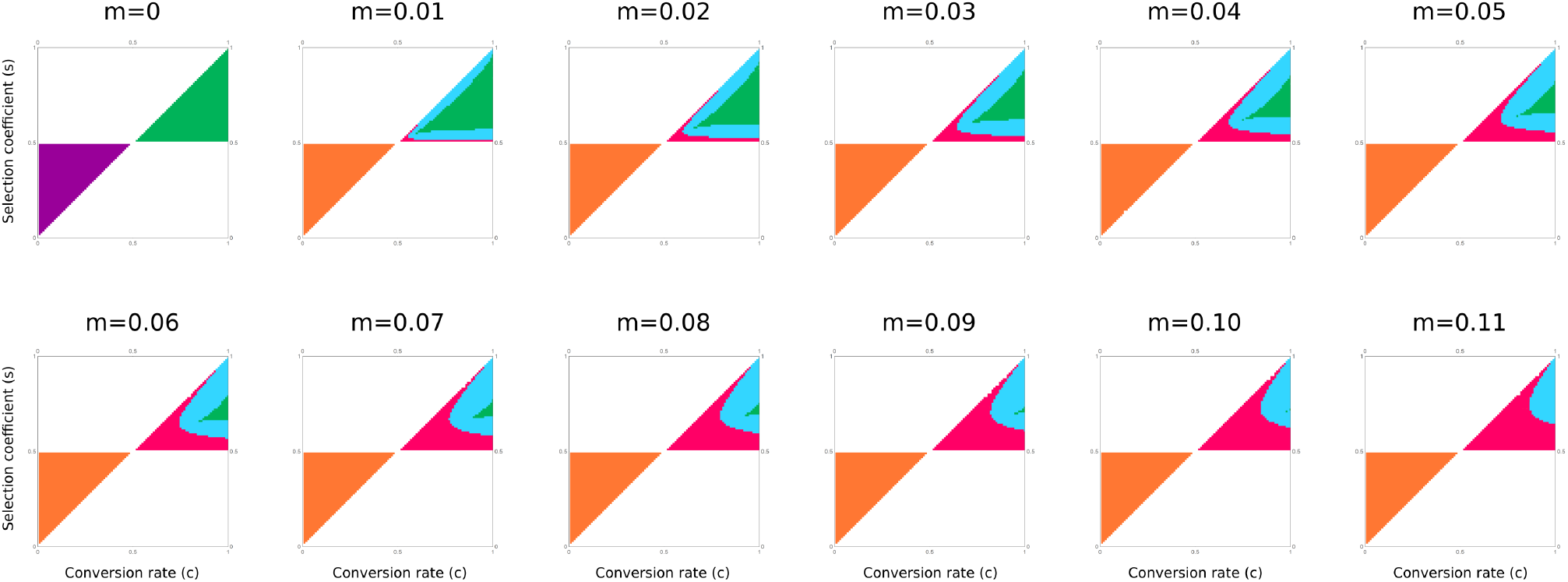
Partitioning of the parameter space by the characterization of the equilibria for the *m*-before-*s* model (Eq. 3), for recessive gene drives (*h* = 0), under different migration rates. Green — 9 equilibria, 4 stable and 5 unstable, one of which is a DTE (stable and 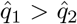); purple — 9 equilibria, 8 unstable and 1 stable and symmetric 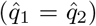; blue — 5 equilibria, 2 stable trivial (global fixation and global loss) and 3 unstable; red — 3 equilibria, 2 stable trivial and 1 unstable; orange – 3 equilibria, 2 unstable trivial and 1 stable symmetric; white — 2 trivial equilibria, 1 stable and 1 unstable. In the bottom white region, the stable equilibrium is global fixation of the gene-drive allele, and in the top white region, the stable equilibrium is global loss. The only gene-drive configurations for which differential targeting is possible appear in green.

**Figure S3:**
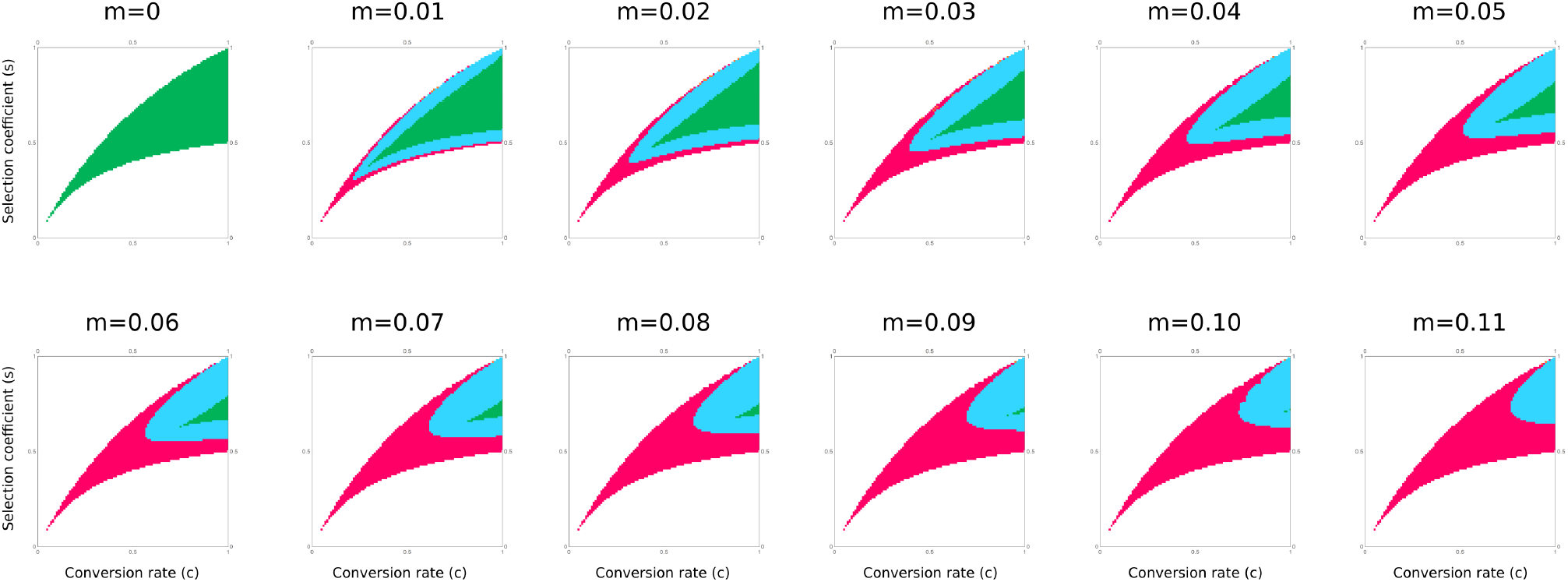
Partitioning of the parameter space by the characterization of the equilibria for the *m*-before-*s* model (Eq. 3), for additive gene drives (*h* = 0.5), under different migration rates. Green — 9 equilibria, 4 stable and 5 unstable, one of which is a DTE (stable and 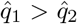); purple — 9 equilibria, 8 unstable and 1 stable and symmetric 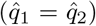; blue — 5 equilibria, 2 stable trivial (global fixation and global loss) and 3 unstable; red — 3 equilibria, 2 stable trivial and 1 unstable; white — 2 trivial equilibria, 1 stable and 1 unstable. In the bottom white region, the stable equilibrium is global fixation of the gene-drive allele, and in the top white region, the stable equilibrium is global loss. The only gene-drive configurations for which differential targeting is possible appear in green.

**Figure S4:**
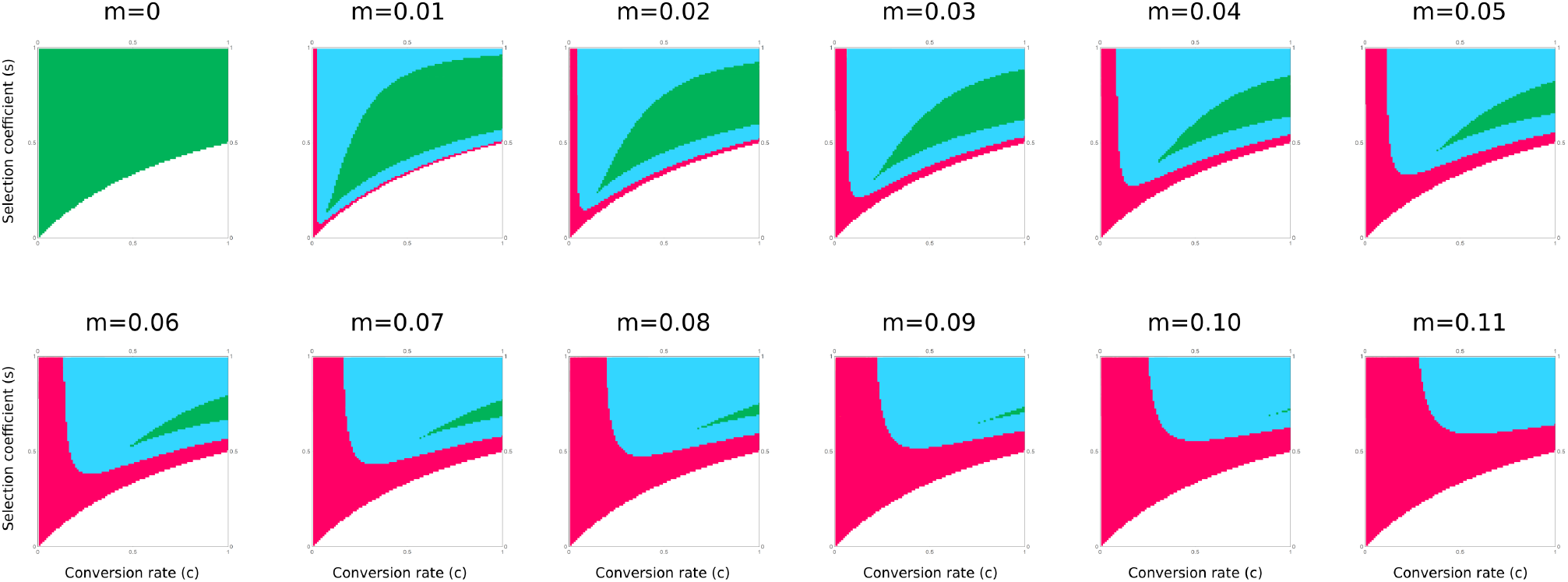
Partitioning of the parameter space by the characterization of the equilibria for the *m*-before-*s* model (Eq. 3), for dominant gene drives (*h* = 1), under different migration rates. Green — 9 equilibria, 4 stable and 5 unstable, one of which is a DTE (stable and 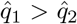); purple — 9 equilibria, 8 unstable and 1 stable and symmetric 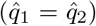; blue — 5 equilibria, 2 stable trivial (global fixation and global loss) and 3 unstable; red — 3 equilibria, 2 stable trivial and 1 unstable; white — 2 trivial equilibria, 1 stable and 1 unstable. In the bottom white region, the stable equilibrium is global fixation of the gene-drive allele. The only gene-drive configurations for which differential targeting is possible appear in green.

**Figure S5:**
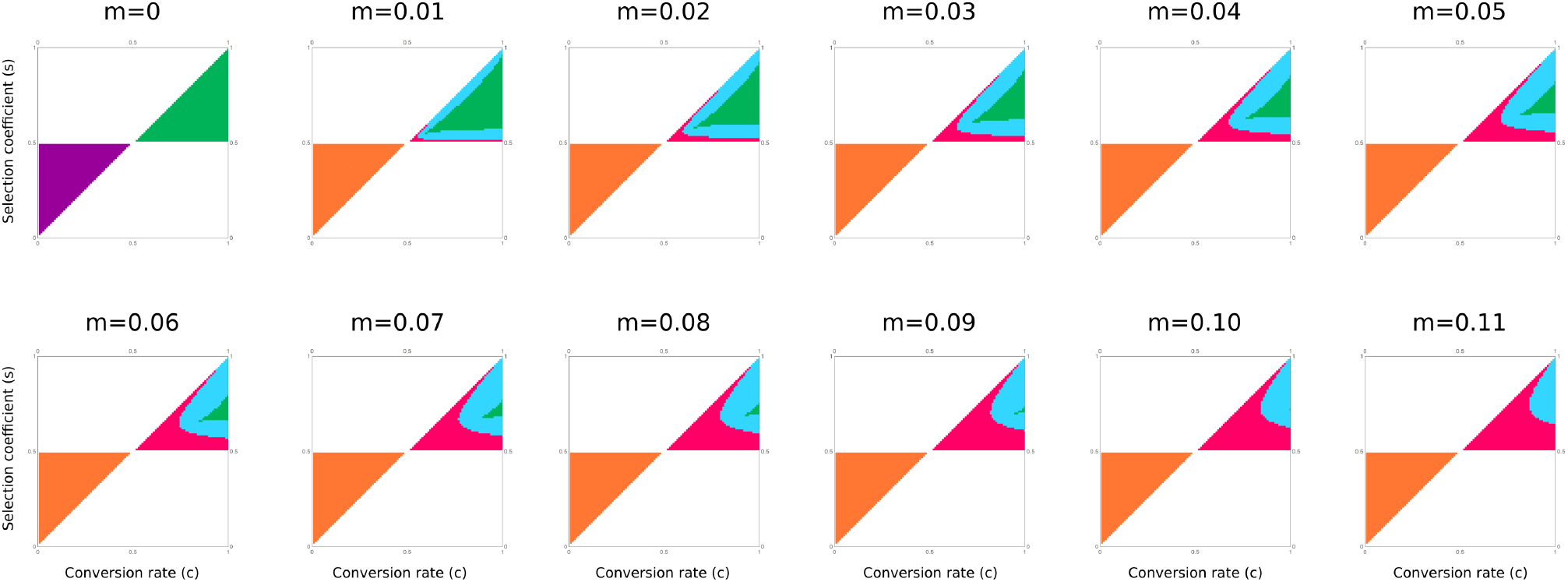
Partitioning of the parameter space by the characterization of the equilibria for the *s*-before-*m* model (Eq. 3), for recessive gene drives (*h* = 0), under different migration rates. Green — 9 equilibria, 4 stable and 5 unstable, one of which is a DTE (stable and 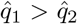); purple — 9 equilibria, 8 unstable and 1 stable and symmetric 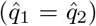; blue — 5 equilibria, 2 stable trivial (global fixation and global loss) and 3 unstable; red — 3 equilibria, 2 stable trivial and 1 unstable; orange – 3 equilibria, 2 unstable trivial and 1 stable symmetric; white — 2 trivial equilibria, 1 stable and 1 unstable. In the bottom white region, the stable equilibrium is global fixation of the gene-drive allele, and in the top white region, the stable equilibrium is global loss. The only gene-drive configurations for which differential targeting is possible appear in green.

**Figure S6:**
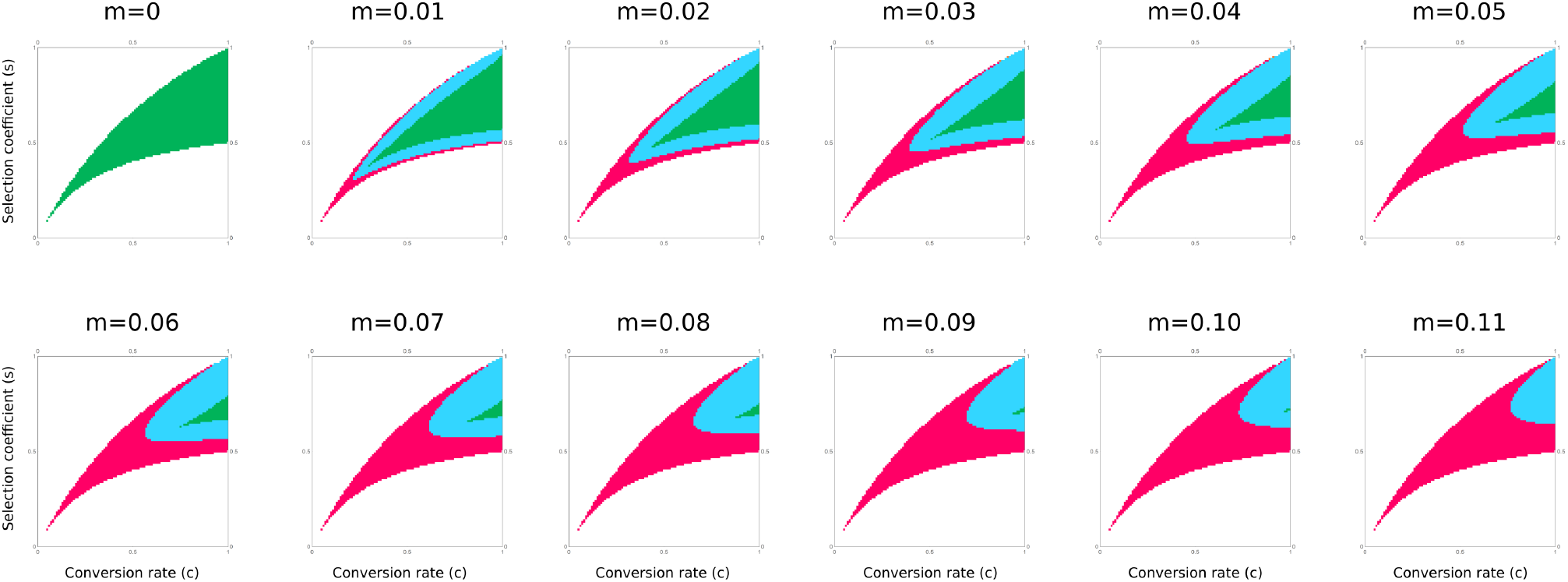
Partitioning of the parameter space by the characterization of the equilibria for the *s*-before-*m* model (Eq. 3), for additive gene drives (*h* = 0.5), under different migration rates. Green — 9 equilibria, 4 stable and 5 unstable, one of which is a DTE (stable and 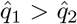); purple — 9 equilibria, 8 unstable and 1 stable and symmetric 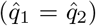; blue — 5 equilibria, 2 stable trivial (global fixation and global loss) and 3 unstable; red — 3 equilibria, 2 stable trivial and 1 unstable; white — 2 trivial equilibria, 1 stable and 1 unstable. In the bottom white region, the stable equilibrium is global fixation of the gene-drive allele, and in the top white region, the stable equilibrium is global loss. The only gene-drive configurations for which differential targeting is possible appear in green.

**Figure S7:**
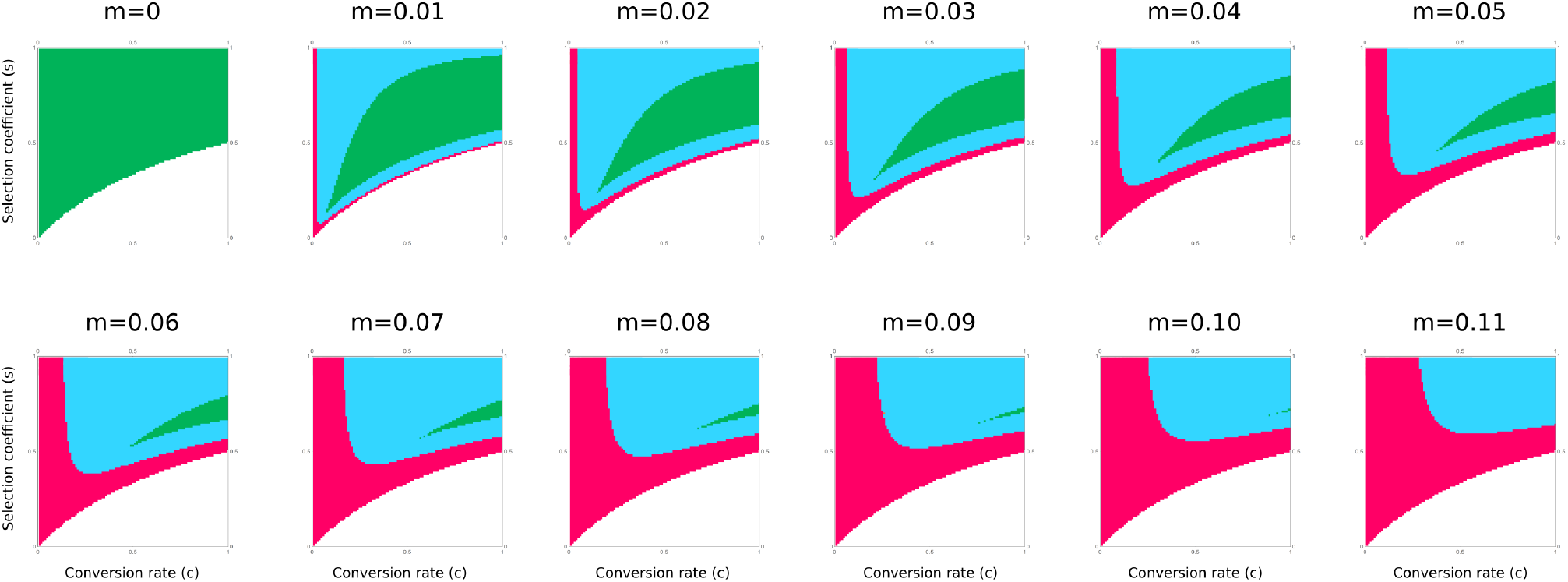
Partitioning of the parameter space by the characterization of the equilibria for the *s*-before-*m* model (Eq. 3), for dominant gene drives (*h* = 1), under different migration rates. Green — 9 equilibria, 4 stable and 5 unstable, one of which is a DTE (stable and 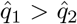); purple — 9 equilibria, 8 unstable and 1 stable and symmetric 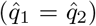; blue — 5 equilibria, 2 stable trivial (global fixation and global loss) and 3 unstable; red — 3 equilibria, 2 stable trivial and 1 unstable; white — 2 trivial equilibria, 1 stable and 1 unstable. In the bottom white region, the stable equilibrium is global fixation of the gene-drive allele. The only gene-drive configurations for which differential targeting is possible appear in green.

